# Task-specific patterns of odorant receptor expression in worker antennae indicates a sensory filter regulating division of labor in ants

**DOI:** 10.1101/2023.01.20.524877

**Authors:** Marcel A. Caminer, Romain Libbrecht, Megha Majoe, David V. Ho, Peter Baumann, Susanne Foitzik

## Abstract

Division of labor (DOL) is a characteristic trait of insect societies, where tasks are generally performed by groups of specialized individuals. In social insects, young workers perform duties within the safety of the nest (e.g., brood care), while older ones undertake riskier tasks (e.g., foraging for food). This DOL remains dynamic, and workers may switch back and forth when colony needs require. Theoretical models propose that workers differ in their thresholds to take on certain tasks when confronted to task-related stimuli, resulting in variation in their response to such stimuli, task-specialization, and thus DOL. Such models assume that workers differ in how they respond to task-related information rather than in how they perceive such information. Here, we test the hypothesis that DOL rather stems from workers differing in their efficiency to detect task-related stimuli. We used transcriptomics to compare gene expression in the antennae and in the brain between nurses and foragers in the ant *Temnothorax longispinosus*. We found that seven times as many genes were differentially expressed between the behavioral phenotypes in the antennae compared to the brain. Moreover, nearly half of all odorant receptors genes were differentially expressed, with an overrepresentation of the 9-exon gene subfamily upregulated in the antennae of nurses. These findings suggest that nurses and foragers differ in how they perceive their olfactory environment, and task-related signals. The results of this study support the hypothesis that a sensory filter in the antennae predisposes workers to specialize in specific tasks, and may improve our understanding of DOL in insect societies.

## Introduction

Division of labor (DOL) is an important organizing principle of complex biological systems that arose independently during three of the major evolutionary transitions [1]. DOL was originally formulated in the context of the production process in human societies [2], but specialization to specific tasks is also found within cells and across many organisms [1]. Examples range from bacteria, where clonal populations are divided into subpopulations focusing on different activities [3–6] to multicellular organisms with differentiation of cells into different tissues and organs, and individuals performing specific roles in animal societies [7,8]. To understand the evolution of complex life it is therefore essential to investigate the mechanisms that underlie DOL.

DOL in insect societies results from individuals specializing in the performance of specific tasks. In addition to the reproductive DOL between fertile queens and functionally sterile workers, there is a behavioral DOL among workers that specialize in tasks such as brood care, foraging, nest building, and defense [9–11]. Several factors can affect task specialization, including age [12,13], nutrition [14,15], morphology [16], genotype [17–19], experience [20], and colony size [21–23]. In most social insect species, workers display an age polyethism by which younger individuals tend to perform intranidal tasks, while older individuals are more likely to undertake activities such as nest-defense and foraging outside the nest [24,25]. Yet, such specialization among workers remains flexible, as foragers can revert to perform brood care when needed [26,27].

Several molecular mechanisms have been implicated in the regulation of DOL. Task specialization is associated with transcriptional changes in the worker brain [28–33]. Molecular pathways such as the insulin/insulin-like signaling (IIS), vitellogenin (Vg) and juvenile hormone (JH) pathways are involved in the regulation of worker behavior [34–39]. Functional manipulations have confirmed that the expression of key genes in the worker brain controls task specialization [40–42]. Behavioral variation among workers is also associated with signaling of biogenic amines (e.g., dopamine, octopamine, tyramine, serotonin), which act as neurotransmitters or neuromodulators involved in the modulation of the responsiveness to task-associated stimuli [43–46].

The self-organization and collective behavior in insect societies are maintained via the exchange of chemical information [47]. Social insects communicate primarily through glandular pheromones and complex mixtures of long-chain hydrocarbons on their cuticle. These cuticular hydrocarbons (CHC) facilitate recognition of nestmates, developmental stages, castes, sexes, and species [48,49]. Social insects perceive chemical information via different types of sensilla on their antennae [50–52]. Decoding the identities of chemical compounds relies on odorant receptors (OR) located within each sensillum [53,54]. OR are transmembrane proteins expressed in the dendrites of olfactory receptor neurons (ORN). The largely conserved OR coreceptor (Orco) of insects [55] is required for odorant recognition in the dendritic membrane: it forms an ion channel with specific OR, which determines the sensitivity and specificity of the ORN [56]. Odorant molecules penetrate through the antenna cuticular pores and are transported by odorant-binding proteins to the ORN membrane, where they interact with receptors, leading to the generation of action potentials [57,58]. ORN axons relay signals from the sensilla to the glomeruli of the antennal lobes in the insect brain, which are the first processing unit for olfactory information. Then, ORN make synaptic contact with the projection neurons and local neurons, which transfer information to the central brain [59–61].

Several lines of evidence indicate that OR genes play central roles in the regulation of social life of insects. First, social insect species typically harbor large numbers of OR genes [62–65]. Second, some OR gene families have specifically expanded during social evolution, such as the 9-exon subfamily in ants, which appears to serve an important function in the perception of CHC [64,66–70]. Third, species that evolved socially parasitic strategies resulting in reduced behavioral repertoires show a strong and convergent reduction in the number of OR genes [71]. Finally, experimentally produced mutant ants that lack the *Orco* gene (coding for the co-receptor necessary for OR to properly function) show impaired social behavior [72,73].

Current DOL models bring together the chemical nature of social insect communication and variation among workers in their response to chemical cues. They posit that flexible response thresholds to task-related chemical stimuli serve as regulators of worker specialization [74]. Workers take on a particular task when the stimulus intensity exceeds their individual threshold for this task. Therefore, individuals with lower thresholds for a given task are more likely to perform it than those with higher thresholds [75]. Individual response decisions and task performance are modulated via numerous parameters on different time scales [7,76–78], and despite extensive research on DOL and task allocation, these mechanisms are not fully understood.

As indicated by the name, “response threshold models” typically assume that individuals differ in their response to specific signals, not necessarily in their ability or efficiency to detect such signals. This implies that such thresholds and the associated responses are set in the central nervous system, possibly via molecular pathways in the brain that correlate with behavioral variation [74,79,80]. We propose that DOL models would benefit from considering odor sensitivity as a potential upstream sensory filter that may affect task specialization. Along these lines, we hypothesize that behavioral variation among workers may also stem from their ability and/or efficiency in detecting different signals. For example, we propose that individuals that specialize in brood care do so because they are better at detecting brood cues, rather than (or in addition to) being more likely to respond to similar levels of brood cues. To test this hypothesis that a sensory filter regulates inter-individual behavioral variation, and thus the DOL in social insects, we investigated transcriptional differences in the brain and antennae of workers that specialize in either brood care or foraging behavior in the ant *Temnothorax longispinosus*. We found that i) behavioral variation was associated with important transcriptomic changes in the antennae, ii) these changes included a large proportion of the OR gene repertoire, and iii) individuals specializing in brood care overexpressed an OR gene family putatively involved in detecting social cues. These findings support our hypothesis that the peripheral nervous system, acting as a sensory filter, plays an important role in regulating behavioral differences between workers and thus in the DOL of social insects.

## Results

### Identification of *Temnothorax longispinosus* behavioral phenotypes

Task specialization in the ant *T. longispinosus* is neither genetically fixed nor rigid, but changes with age and in response to colony needs [38]. We conducted behavioral observations of seven *T. longispinosus* laboratory colonies to identify individuals that specialize in brood care behavior (hereafter referred to as nurses), and others that specialize in foraging (hereafter referred to as foragers) (Fig 1*A*; see Methods section for more details). Individuals identified as nurses interacted with the brood in 54% ± 22% (mean ± sd) of the observations, and were recorded outside the nest in 1% ± 3% of the observations. On the contrary, foragers were found outside the nest in 42% ± 26% of the observations, but interacted with the brood in only 2% ± 4% of the observations.

**Fig 1.**
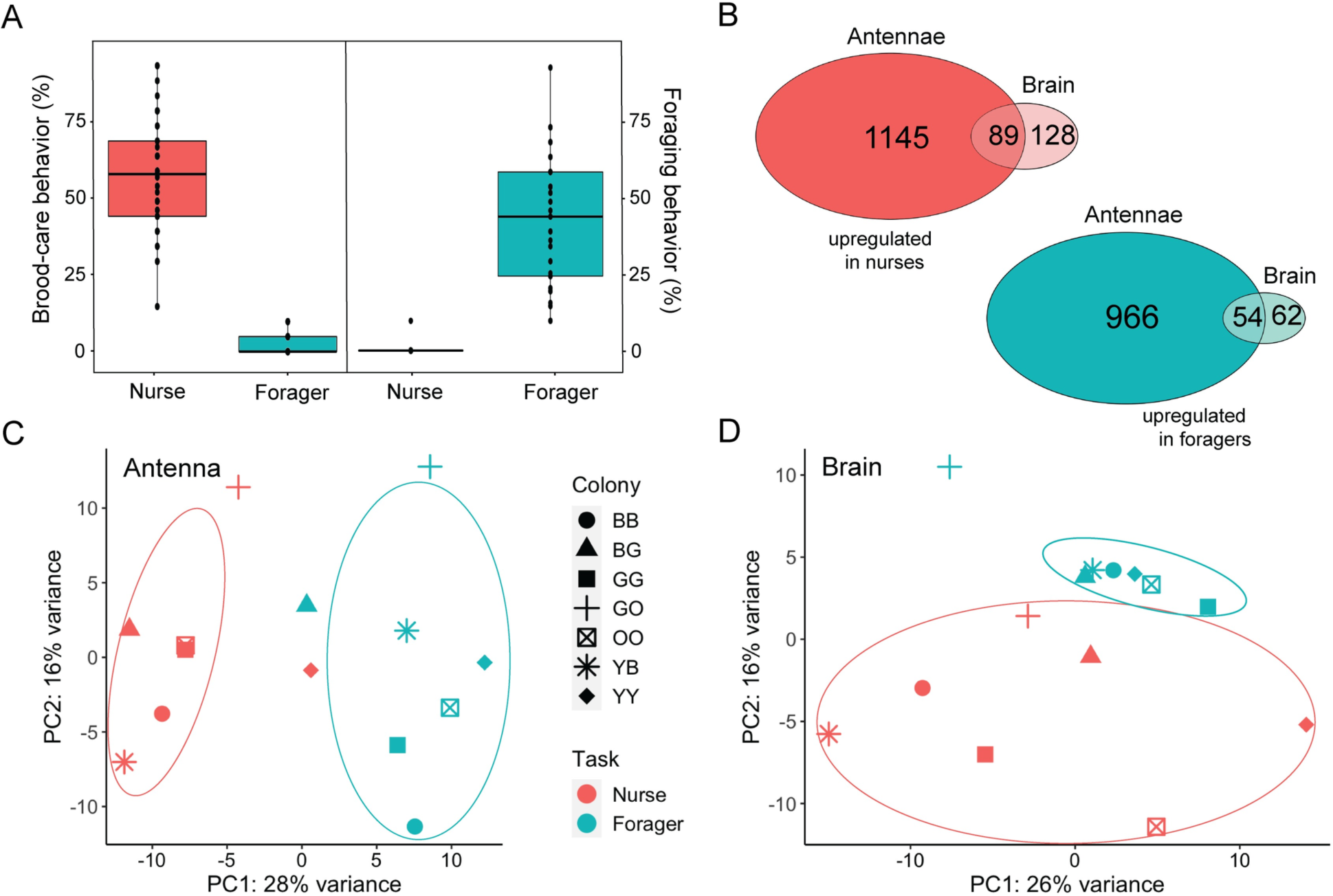
Variation in behavior, brain and antenna gene expression between nurses and foragers. (*A*) Boxplot showing behavioral differences between ants selected for transcriptomic analysis. Each black dots represent individual ants. For differential expression analysis, each sample contains the pooled RNA of seven ants of the respective behavioral phenotype. (*B*) Venn diagrams showing the number of DEG that were overexpressed in nurses (red) and foragers (turquoise) for both antenna and brain tissues. Principal component analysis (PCA) plots based on all expressed genes for (*C*) antenna and (*D*) brain samples. The color of each sample represents task (red = nurse, turquoise = forager), and the shape the colony of origin. The ellipses were defined by a 95% confidence interval and show how similar/different the samples are within task groups.

### Larger task-associated transcriptomic changes in the antennae than in the brain

To investigate transcriptomic variation between nurses and foragers, we used RNA-seq to generate seven nurse and seven forager brain and antenna samples, each consisting of pooled tissue from seven workers of a single colony. Of the 14,837 protein-coding genes annotated in the *T. longispinosus* genome, we found that 91% (13,494) and 92% (13,683) were expressed (FPKM > 0) in the brain and the antennae, respectively. Using an adjusted p-value of 0.05 as significance threshold, we detected 333 differentially expressed genes (DEG) in the brain (217 upregulated in nurses, 116 in foragers), and 2,254 in the antennae (1,234 upregulated in nurses, 1,020 in foragers; Dataset S1; Fig 1*B*). We found an overlap of 162 DEG between the two tissues, including 143 DEG that showed differences in the same direction across tissues. Principal component analysis (PCA) plots indicate that both brain and antenna samples cluster by behavioral phenotype rather than the colony of origin (Fig 1 *C*-*D*).

### OR gene expression differs between antennae of nurses and foragers

To investigate whether odorant perception differs between nurses and foragers, we focused our attention on the expression of OR genes in the antennae. We found that all 419 OR genes in the genome of *T. longispinosus* [71] were expressed in the antennae, and that 48% (202/419) of them were differentially expressed between nurses and foragers. Specifically, 65 OR genes were upregulated in nurses (15% of all OR genes), and 137 in foragers (32% of all ORs) (Fig 2). Then, we studied which OR subfamilies were preferentially expressed in nurses and foragers. The 65 OR genes overexpressed in nurses belonged to three OR subfamilies, while the 137 OR genes upregulated in foragers were distributed among 19 OR subfamilies. Foragers overexpressed 26, 8, and 8 OR genes from the L, P, and H subfamilies, respectively, while no OR genes from these subfamilies were upregulated in nurses (Dataset S2). We found that 61% (26/43) of the genes from the L subfamily were overexpressed in foragers, which represents a significant overrepresentation (Fisher’s test, odds ratio = 3.63, p-value < 0.001). For the P and H subfamilies we did not find such an overrepresentation, likely due to lower gene numbers (P subfamily: Fisher’s test, odds ratio = 3.62, p-value = 0.06; H subfamily: Fisher’s test, odds ratio = 1.87, p-value = 0.19). On the other hand, we found that 82% (53/65) of the OR genes overexpressed in nurses belong to the 9-exon subfamily. This results in an overrepresentation of the 9-exon subfamily in genes that were overexpressed in nurses (Fisher’s test, odds ratio = 20.98, p-value < 0.001), with 46% (53/114) of this subfamily being overexpressed in nurses. In contrast, only 6% (8/137) of the 9-exon subfamily was overexpressed in foragers, which is less than expected by chance (Fisher’s test, odds ratio = 0.10, p-value < 0.001). We also found that nurses overexpressed *Orco* compared to foragers (FDR p-value = 0.001).

**Fig 2.**
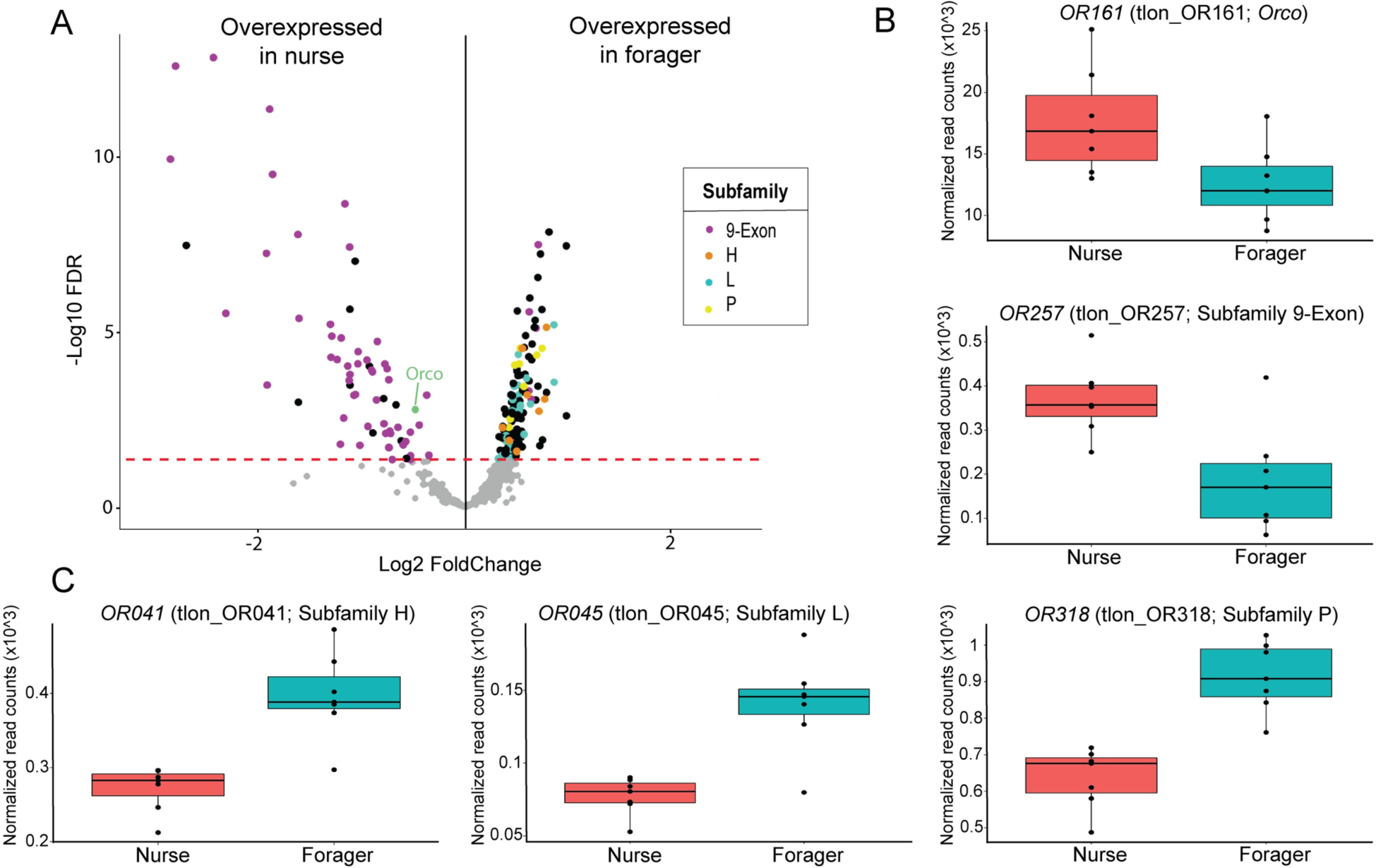
OR expression in the antennae differs between nurses and foragers. (*A*) Among the DEG overexpressed in nurses and foragers, the most represented OR subfamilies are indicated by different colors. The red dotted line represents the significance threshold of our differential expression analysis. (*B*,*C*). Boxplots representing the expression of (*B*) *Orco* and *OR257*, the gene with the highest expression difference in terms of log2 FoldChange in the 9-exon subfamily, and (*C*) *OR041*, *OR045* and *OR318*, the genes with the highest expression difference in terms of log2 FoldChange in the H, L and P subfamilies, respectively.

### Genes in biogenic amine pathways vary in expression between nurses and foragers

Because biogenic amines are implicated in the regulation of behavior, and they may affect sensory perception [45,81–83], we screened our lists of DEG in the brain and antennae for genes in biogenic amine pathways. We found five genes associated with biogenic amine signaling in the antennae, and none in the brain. Genes in the serotonin (*5- hydroxytryptamine*, DBV15_11483), tyramine (*tyramine beta-hydroxylase*, DBV15_00422) and octopamine (*octopamine receptor* LOC112465659) pathways were upregulated in foragers, while nurses showed a higher expression of genes in the dopamine (*dopamine 1-like receptor 2*, DBV15_07611) and octopamine (*octopamine beta2 receptor*, DBV15_10418) pathways (Fig 3 *A-E*).

**Figure 3.**
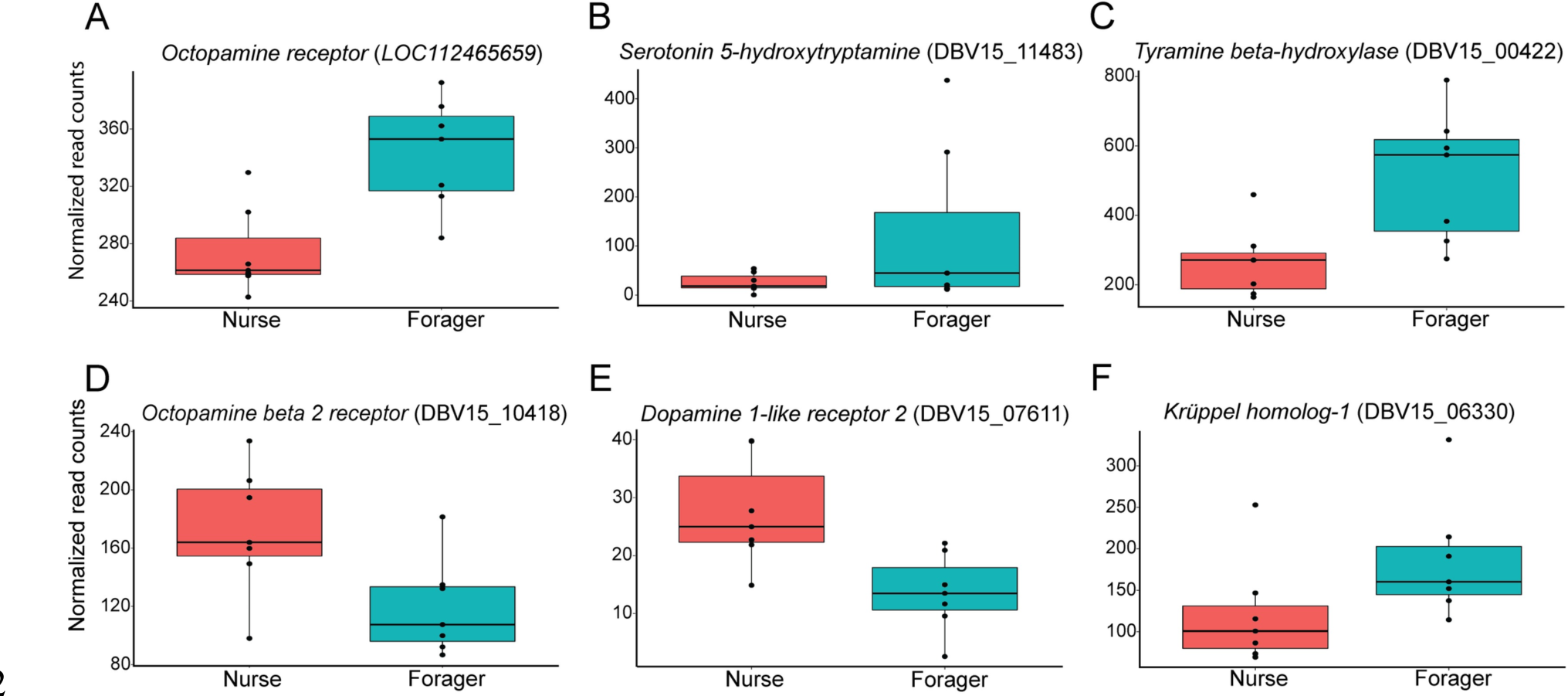
Gene expression differences between nurse and forager antennae. Boxplots representing the expression of (*A*-*E*) selected genes in biogenic amines pathways, as well as (*F*) one behavioral candidate gene (Table S1). Gene identity is shown in parentheses.

### Association between behavioral variation and molecular pathways in brain and antennae

Our analysis revealed expression differences between the brains of nurses and foragers for genes involved in the regulation of task specialization in social insects. Previous studies have found that *Vg* genes and the associated JH and IIS/TOR pathways are important endocrine networks that play central roles in the regulation of lifespan, fertility and behavior in bees and ants [35,84–89]. Therefore, we searched our RNA-seq data for task-associated expression of genes involved in the metabolism, biosynthesis, and regulation of these pathways in the brain and antennae. We found evidence that multiple genes from these pathways are differentially expressed between nurses and foragers (Table S1). Among the most interesting genes upregulated in the brain of nurses were *conventional Vg* (LOC112466671), *Vg-like A* (named/classified as per [42]; DBV15_03138), *venom carboxylesterase-6-like* (DBV15_11528), and *protein takeout* (DBV15_08771), while *allatostatin A-like* (LOC112454443) and *insulin-like growth factor I* (*IGF1*) (LOC112454447) were overexpressed in the brain of foragers. Remarkably, *venom carboxylesterase-6-like* and *IGF1* were overexpressed in the antennae of nurses and foragers respectively. Additionally, high expression levels of the zinc-finger transcription factor *Krüppel homolog-1* (*Kr-h1*) were detected in the antennae of foragers (DBV15_06330) (Fig 3*F*). The expression of this gene has been correlated with caste and behavioral differences in the brain of social insects [86,90–93], but little is known about its function and gene targets in the antennae.

We performed a GO enrichment analysis to gain a deeper understanding of the biological processes represented in the lists of DEG. This analysis detected many enrichments based on single one or few genes, and we mention below a few interesting processes with the number of genes driving the enrichment in parentheses (see Table S2 for complete list). Genes that were upregulated in the brain of nurses were enriched for biological processes such as *translation* (19 genes), *cellular iron ion homeostasis* (2 genes), and *catabolic process* (6 genes); while *translation* (22 genes), *endocytosis* (6 genes), *glucose metabolic process* (4 genes), and *regulation of cell cycle* (3 genes) were enriched in the antennae. In the list of genes that were overexpressed in the brain of foragers, we found that enriched processes included *peptide metabolic process* (4 genes), *methionyl-tRNA aminoacylation* (1 gene), *positive regulation of type I interferon production* (1 gene), and *cell redox homeostasis* (1 gene); while *phosphatidylinositol phosphorylation* (4 genes), *inositol phosphate dephosphorylation* (3 genes), *carbohydrate metabolic process* (17 genes), *innate immune response* (2 genes), and *nucleotide catabolic process* (3 genes) were enriched in the antennae.

## Discussion

In this study, we tested the hypothesis that variation among social insect workers in their ability to detect different chemical signals regulates task specialization, and thus DOL in insect societies. To do so, we compared the brain and antenna transcriptomes of nurses and foragers in the ant *T. longispinosus.* We report several lines of evidence that support our hypothesis. First, we found almost seven times as many genes to be differentially expressed between nurses and foragers in the antennae as in the brain, indicating that peripheral sensory organs may be more important than the central nervous system in the task specialization process of social insect workers. Second, we found half the OR genes in the *T. longispinosus* genomes are differentially expressed between nurse and forager antennae, suggesting that behavioral specialization is associated with different sensory filters that result in specific perceptions of the chemical environment. Third, our analyses revealed that nurses and foragers upregulated distinct families of OR genes, indicating that their sensory filters may target different types of chemical cues, possibly adjusted to the tasks they perform. Finally, we detected several genes in multiple biogenic pathways to be differentially expressed between the brains of nurses and foragers, potentially involved in fine-tuning the sensitivity of the odorant filters.

Many organisms have evolved sensory filters to focus on only a subset of all environmental cues. In insects, the peripheral olfactory filtering system enables individuals to detect and discriminate odors that convey ecologically relevant information used to mediate important behaviors such as courtship, locomotion and navigation to avoid predators and locate food or nesting sites [94]. For example, the mosquito *Anopheles gambiae* strongly responds to odorant components of its vertebrate hosts that provide meals [95]. Sensory filters have been selected for because they limit the amount of information perceived by the organism, and thus the energy and time required by the brain to process it. Within insect colonies, individuals typically exhibit many morphological and physiological traits associated with increased efficiency in task specialization. They can be understood as adaptations that allow individuals to become more proficient at their task due to learning, training, or the perceiving valuable task-related information, allowing the colony to avoid the cost of task switching [96]. In this context, it is interesting to find that *T. longispinosus* workers may have different sensory filters that would serve as a basis for their task specialization, as workers would mostly perceive chemical cues that pertain to their tasks. Our findings indicate that the sensory filter of ant workers is dynamic, and its changes may underlie their behavioral maturation.

Age, genetic background, social environment, and individual experience produce behavioral variation among workers that results in division of labor [13,20,23,97]. In our study, age in particular likely differed between nurses and foragers. However, previous experiments designed to disentangle gene expression associated with behavioral specialization, age and fertility showed that behavioral specialization is more strongly associated with gene expression than age and fertility in *T. longispinosus* [38]. We propose that the factors known to regulate task specialization could drive different OR expression patterns, which in turn would produce behavioral variation via contrasting abilities to detect different sets of odors. This hypothesis is supported by several studies (reviewed in [98]) showing that the physiological condition of an animal can influence the level of receptor expression, including mating status, oviposition, feeding, circadian rhythm, experience, and aging. Alternatively, we cannot exclude that the exposure to different odors that is associated with performing different tasks may at least in part have affected OR gene expression in the antennae. However, such an effect is unlikely to explain the large-scale variation in gene expression, as there is very limited evidence that the mere exposure to odors influences the expression of the gene coding for the OR that binds this odor [99,100].

According to our sensory filter hypothesis, ant workers would be expected to primarily detect chemical cues that correspond to their tasks. We found that the 9-exon subfamily of OR was overrepresented in genes that were upregulated in the antennae of nurses. Interestingly, recent studies have shown a rapid expansion of the 9-exon subfamily in ants, and several lines of evidence indicate that OR genes from this subfamily mediate complex social interactions in ant colonies [65–67,69]. First, comparisons of antennal transcriptomes revealed that 9-exon OR genes are expressed more frequently in workers than in males, suggesting a role in social communication among workers [68,69]. Second, representative OR from the 9-exon subfamily can detect CHC extracts from several castes [70]. According to McKenzie et al. [69], 9-exon OR genes were first expressed in solitary ancestors of aculeate wasps and facilitated CHC discrimination, likely for prey or mate recognition, with a lineage giving rise to the ancestors of ants. Moreover, OR genes of the 9-exon subfamily were convergently lost in socially parasitic ant species that lost the ability to perform brood care or foraging [71], suggesting that they are essential for the performance of these worker tasks. Our finding of an overexpression of 9-exon OR genes in the antennae of nurses is in line with these previous studies, and suggests that many of these receptors have important functions within the nest, such as sensing chemical cues from the larvae, queen, or other workers, and/or that they are less important for sensing task-related stimuli or other signals outside the nest. Among the OR genes overexpressed in nurses was also the co-receptor *Orco*, which is widely expressed in olfactory sensory neurons and nearly unchanged in sequence in distant insect taxa [101–103]. The overexpression of *Orco* may indicate a generally more active odorant sensory system in the antennae of nurses compared to foragers.

The behavioral transition from nursing to foraging may be triggered by a lower efficiency in detecting brood cues via the downregulation of specific OR genes (e.g., from the 9-exon subfamily). This would result in ants moving farther away from the brood, and this change in spatial location may trigger the behavioral transition to outside tasks [104,105]. In addition to being less efficient at detecting brood cues, the sensory filter of foragers may also become fine-tuned to detect a more diverse set of odors. Foragers overexpressed a greater number of OR gene subfamilies compared to nurses (19 and 3 for foragers and nurses, respectively), which may indicate that the olfactory system of foragers could be adjusted to the more diverse chemical environment outside the nest. Similar to the 9-exon subfamily, the L subfamily has also been expanded in social insects [62,64,65], and along with the P and H subfamilies, it has been lost in socially parasitic ants [71]. Interestingly, the OR genes from the L, P, and H subfamilies have been upregulated in foragers, and thus may have a task-specific function, such as recognition of chemical cues related to environmental perception or recruitment cues outside the nest. To date, however, there is no information on which odors are recognized by these OR gene subfamilies.

Finding task-specific variation in OR gene expression raises the question as to which molecular mechanisms regulate those changes. Previous studies in insects revealed that olfaction-guided behavior is mediated by biogenic amine receptors in the antenna, and their expression is involved in fine-tuning the sensitivity of the olfactory system [82,98]. For example, modified concentration cAMP and intracellular Ca^2+^ levels due to octopamine-induced signal transduction in the moth *Manduca sexta* [106] activate Orco, leading to changes in ORN sensitivity [107,108]. Our results suggest that the biogenic amine signaling pathway may modulate the sensory filtering function of insect antennae and alter sensitivity to various signals. We found that genes encoding tyramine and its precursor, octopamine, are upregulated in forager antennae, similar to genes involved in serotonin signaling. Tyramine and dopamine (which was upregulated in nurses) have been implicated in modulating taste and olfactory receptor neurons, while serotonin may serve as a neurotransmitter and neurohormone in antennal vessels and mechanosensory organs [82]. Serotonin influences foraging activity [81] and regulates food intake in many animals [109–111]. Dopamine signaling also plays an important role in controlling the insect circadian clock and mediating clock-controlled behavioral phenotypes such as locomotion [112,113]. In our focal species *T. longispinosus*, inside workers were found to exhibit a stronger circadian rhythmicity than foragers, which may be regulated via differences in the acetylation of histone proteins [114].

Changes in behavior and olfactory sensitivity in insects could be related to the expression of genes involved in IIS, target of rapamycin (TOR), JH and Vg pathways, according to age, circadian rhythm, mating and feeding status [98]. For example, appetite state in *D. melanogaster* is signaled by insulin, which upregulates a peptide receptor on the olfactory receptor cells that innervate the DM1 glomerulus. Activation of the DM1 glomerulus is enough to drive the fly to reach for food [115]. Recent studies have shown that pheromone release and odor sensitivity appear to be under JH control in *Schistocerca gregaria* and *Locusta migratoria*, which could lead to behavioral changes [116–120]. Finally, an experimental downregulation of *Vg-like A* in *T. longispinosus* workers resulted in decreased brood care behavior and a lower sensitivity to brood-related chemical cues, suggesting changes in odor perception and olfactory-driven decision making [42]. Our results reveal that genes associated with all these pathways were differentially expressed in the brain and antennae between nurses and foragers, predicting a link between their role in task-associated behavioral changes and the regulation of odor perception. Given the central role of IIS, Vg, JH, and TOR pathways in regulating division of labor in social insects, and our finding of task-associated patterns of the antennal expression of genes from multiple biogenic amines, we hypothesize that these modulators and hormones could be involved in the regulation of the olfactory filter. How the detailed molecular mechanisms of these pathways in the brain are causally linked to the complex changes in olfactory perception in the antennae should therefore be investigated next.

## Conclusion

Our transcriptomic analyses of the brain and antennae of *T. longispinosus* nurses and foragers provide support to our hypothesis that behavioral variation and task specialization in ant workers are regulated via differences in olfactory perception. We predict that antennal physiology acts as sensory filters that limit the type and amount of chemical information passed to the brain. This would allow workers to target relevant chemical information from the environment and discriminate signal from noise without using energetically costly processing by the central nervous system. We argue that this sensory filter is flexible and regulated through changes in physiological conditions such as age, nutrition, and hormones. Variation among workers in their efficiency to detect specific chemical cues would result in task specialization and division of labor. Our study opens novel avenues of research to better understand the role of sensory filters in controlling DOL in insect societies.

## Materials and Methods

### Sample collection and behavioral determination

A total of seven colonies of the ant *T. longispinosus* were selected with an average colony size of 110 ± 31.5 workers (mean ± SD, Dataset S3). The ants were collected in the forests of the Edmund Niles Huyck Preserve, Renssellearville, NY, USA (42°31′41.0′′N 74°09′38.8′′W), in June of 2018 with permission. Upon collection, we housed each colony in a plaster-floored nesting box (43 cm × 28 cm × 10 cm) divided into three chambers containing a single slide nest, in which the colony relocated. A slide nest is an artificial nesting site comprised of a small Plexiglas cavity sandwiched between two glass microscope slides. Colonies were established at the Johannes Gutenberg University in Mainz, Germany, under a 14 h:10 h light:dark photoperiod at 18°C to a 22°C temperature. We provided honey and water ad libitum and fed crickets to the colony twice a week. To allow for visible behavioral division of labor between workers of the two behavioral phenotypes, we marked, observed and recaptured ants from inside and outside the nest. We defined foragers as workers that perform outside-nest tasks, including foraging for food, while nurses remained inside the dark nest and cared for the ant brood. A total of 69 workers inside (from the brood pile) and 76 workers outside the nest were marked with fine colored metal wires (0.02 mm Elektrisola, Eckenhagen, Germany) between the petiole and post-petiole. We performed behavioral observations every two hours, four times a day for five days (total = 20 scans), in which we noted down how many times an individual spent performing brood care and foraging behavior and the position in the nest (Table S3). Based on these behavioral observations, the marked individuals found outside the nest, exploring the surroundings for food or water, were identified as foragers.

These workers usually do not care for the brood and do not frequently reside on brood piles, as ant colonies organize themselves spatially in a way that reduces contact between foragers and brood [121]. We identified nurses as workers that remain inside the nest in direct contact with brood and were unlikely to leave the nest. Foragers were found in 42% ± SD 26% of the observations outside of the nest, whereas nurses spend only 1%± SD 3% outside. In contrast, nurses were interacting with the brood in 54% ± SD 22% of the observations, whereas we found that foragers only did this only in 2% ± SD 4% of the observations. We scanned the behavior of workers over 20 observations, albeit earlier studies have shown that a single observation allows to group *T. longispinosus* and other ants reliably into nurses and foragers that differ in behavior [121,122], gene expression [38] and CHC composition [42]. Furthermore, spatial location can alone can predict behavior in *Temnothorax* workers [123]. We focused in this study on individuals highly specialized on either foraging or brood care. Workers that performed both tasks regularly were not included in this study. After all observations were completed, the marked nurses and foragers were collected, directly frozen in liquid nitrogen and stored at -80 °C until further processing for dissection and pooling according to behavioral state and colony.

### RNA extraction and sequencing

For RNA extraction, we removed both antennae and stored them in a 1.5 ml Eppendorf tube containing 50 μl TRIzol (Invitrogen), cut the head off and fixed it on a slide with melted dental wax. We then made an incision around the head with a surgical scalpel and removed the head capsule with forceps to expose the intact brain. Finally, we carefully pulled the brain out of the head capsule and removed the remains of other tissues that were connected to it. The dissected brain was transferred to a 1.5 ml Eppendorf tube containing 20μl PBS. Each dissection was completed in less than 5 minutes to prevent RNA degradation. We dissected brain and antennae tissues from 48 nurses and 49 foragers. We pooled the brains and antennae from seven workers from each behavioral state and colony. The only exception was the “GO” colony (NY18 E110), for which we pooled only six brains and twelve antennae from six nurses (Dataset S3) due to the loss of one sample during the dissection process. Immediately after dissection of each brain and antennae, the Eppendorf tubes were kept on dry ice while we dissected the remaining individuals. Brain and antennae tissues were homogenized with a pestle. Sample brains were transferred separately to a 1.5 ml Eppendorf tube containing 50μl of TRIzol. We added 50μl chloroform to each brain and antenna samples, cautiously inverted for 30 s and then centrifuged samples at 12,000 g for 15 min at 4°C. We collected the resulting supernatant and precipitated RNA with 25μl 70% ethanol. We conducted the subsequent RNA extraction with the RNeasy Mini Kit (Qiagen), following the manufacturer’s instruction. The resulting 28 samples (14 brains and 14 antennae) were stored at -80°C until library preparation.

RNA-seq libraries were prepared by Novogene Company Limited, Cambridge, UK, using the NEBNext Ultra RNA Library Prep Kit for Illumina according to the manufacturer’s protocol. After amplification and purification, 28 libraries were sequenced on an Illumina NovaSeq 6000 S4 flow cell platform using a paired-end 150 bp. Approximately 43 million raw reads were generated from each library (Dataset S3).

### Gene expression analyses

Raw data obtained from Novogene were checked using FastQC v.0.11.9 [124], and Illumina adapters were removed using Trimmomatic v.0.36 [125]. The protein-coding genes of *T. longispinosus* (GCA_004794745.1; [126] and the congener *T. curvispinosus* (GCA_003070985.1) were retrieved from the NCBI database and transferred to the recently published *T. longispinosus* genome [71] using the Liftoff v.1.6.1 tool [127]. In total, 10,029 of 13,061 (∼77%) annotated protein-coding genes were transferred from the original *T. longispinosus* assembly (genes identified as “DBV15”) and 4,808 were transferred from *T. curvispinosus* (genes identified as “LOC”), for a total of 14,837. For gene expression analysis, reads were mapped to our *T. longispinosus* genome assembly, and the read counts table was generated using STAR 2.7.0 [128] with default settings. Detailed mapping statistics for each sample is available in Dataset S3. We used the deseq2 v1.16.1 package to identify differentially expressed genes [129]. To avoid biased results due to low read counts, we removed from the counting matrix those genes for which less than 10 of the reads mapped to at least 6 of our 14 samples (n - 1 of the smallest sample size). Then, we conducted a differential gene expression analysis with DESeq2 [130]. We began with comparisons between nurses and foragers using the ∼Colony+Task model, followed by a likelihood ratio test (LRT) approach, with colony ID as a fixed factor. Genes were considered differentially expressed if the false discovery rate (FDR), using Benjamini-Hochberg procedure, had an adjusted p-value of ≤ 0.05. The resulting lists of DEG refer to genes that are overexpressed and underexpressed in foragers compared to nurses. We used the online tool Venny v.2.1 (https://bioinfogp.cnb.csic.es/tools/venny) to generate a Venn diagram containing the DEG associated with task and tissues. Separation of differentially expressed genes by task was visualized by performing principal component analysis (PCA) on transformed reads of filtered transcriptomes from all contigs using the plotPCA function provided by DESeq2. Finally, ORs genes that were upregulated in each behavioral phenotype was visualized in a volcano plot using the EnhancedVolcano package in R.

### Identification of behavior candidate genes and odorant receptors

We used gene annotations based on a BlastX search of the *T. longispinosus* transcriptome compared to a list of different invertebrate proteomes (i.e., *Acromyrmex echinatior, Apis mellifera, Camponotus floridanus, Drosophila melanogaster, Harpegnathos saltator, Odontomachus brunneus, Temnothorax curvispinosus*) downloaded from the NCBI database with an E-value of 1e-5 and below. Clusters containing more than one sequence match per species were reduced to a single specimen based on the highest blast score. We constructed orthogroups across all of the above species using OrthoFinder [131], including amino acid sequences from the *T. longispinosus* proteome [126], and retained orthogroups containing caste DEG (Dataset S4) to again compare potential behavioral candidate genes previously identified as involved in regulating the division of labor in social insects [38,41,87,93,120,132–135]. GO enrichment analysis was performed with TopGO v.2.44.0 using a Fisher’s exact test for the different gene sets compared to the whole genome with the weight01 algorithm [136]. Only annotated GO terms with an adjusted p-value of ≤ 0.05 were considered significantly enriched. All enrichment analyses were performed in RStudio v.1.4.1106 [137].

Odorant receptor (OR) protein sets were clustered across multiple ant species using OrthoFinder to derive orthologous groups and identify subfamilies for each OR in *T. longispinosus*. To associate orthogroups with previously identified OR subfamilies, we used OR annotation in *Atta cephalotes, Acromyrmex echinatior* from Engsontia et al. [65], and *Camponotus floridanus*, *Harpegnathos saltator*, and *Solenopsis invicta* from Zhou et al. [64,68]. Missing subfamily information was labeled as “unassigned” (Dataset S2).

## Acknowledgments

We thank Diego Páez-Moscoso for many helpful discussions and comments throughout the study; Marah Stoldt and Carlotta Martelli for providing helpful comments on the manuscript; and Marion Kever and Jenny Fuchs for technical assistance. This project was funded by the Deutsche Forschungsgemeinschaft (DFG, German Research Foundation) – GRK2526/1 – Projectnr. 407023052.

## Data

RNAseq data that support the findings of this study have been deposited in GenBank with the BioProject accession codes PRJNA926589 (http://www.ncbi.nlm.nih.gov/bioproject/926589).

## Supporting Information

Table S1. List of candidate genes related to behavior phenotypes.

Table S2. List of enriched GO biological process terms of the DEG in brain and antenna.

Table S3. Behaviors and positions annotations during nest scans. Nursing was defined as the number of observations a worker antennating, grooming, feeding or carrying a brood item, and its position inside the nest (on the brood or near to the brood pile). Foraging was defined as the number of observations an individual was found outside the nest, collecting food or water.

Dataset S1. Gene list with caste-differential expression, NCBI BlastX results, and gene read count of 28 RNA-seq samples.

Dataset S2. OR subfamilies and orthogroups from the DEG lists between nurses and foragers in the antenna.

Dataset S3. Summary of ant brain and antenna transcriptome data sets and mapping results.

Dataset S4. List of orthogroups from the DEG list between nurses and foragers in *Temnothorax longispinosus*.

