## Supplemental Table S1 for "Task-specific patterns of odorant receptor expression in worker antennae indicates a sensory filter regulating division of labor in ants"

**Table S1. List of candidate genes related to behavior phenotypes.**

| Gene Annotation | Locus (*T. longispinosus*) | Organism reference | Locus reference | Orthogroup | Task | Tissue | Pathways and genes | References |
|---|---|---|---|---|---|---|---|---|
| protein takeout isoform | DBV15_08771 | *Camponotus floridanus* | LOC105257386 | OG0007297 | Nurse | Brain | JH: signalling | [1] |
| vitellogenin-1 (Vg conventional) | LOC112466671 | *Harpegnathos saltator* | LOC105181726 | OG0003549 | Nurse | Brain | Vitellogenin | [1–4] |
| vitellogenin-2-like (Vg like-A) | DBV15_03138 | *H. saltator* | LOC105191299 | OG0009030 | Nurse | Brain | Vitellogenin | [2,3] |
| farnesyl pyrophosphate synthase | LOC112466955 | *Drosophila melanogaster* | Fpps | OG0000401 | Nurse | Brain | JH: biosynthesis | [5] |
| venom carboxylesterase-6 (Jhe1) | DBV15_11528 | *H. saltator* | LOC112590013 | OG0002262 | Nurse | Brain/Antenna | JH: regulation of JH biosynthesis | [1,3] |
| ribosomal protein S6 | DBV15_00490 | *D. melanogaster* | RpS6 | OG0007546 | Nurse | Brain/Antenna | IIS / TOR | [5] |
| dopamine 1-like receptor 2 | DBV15_07611 | *D. melanogaster* | Dop1R2 | OG0001377 | Nurse | Antenna | bioagenic amines | [6] |
| octopamine beta2 receptor | DBV15_10418 | *D. melanogaster* | Octbeta2R | OG0000466 | Nurse | Antenna | bioagenic amines | [6] |
| aldehyde dehydrogenase | DBV15_01763 | *D. melanogaster* | Aldh-III | OG0000175 | Nurse | Antenna | JH: biosynthesis | [5] |

| | | | | | | | | |
|---|---|---|---|---|---|---|---|---|
| cyclin G | DBV15_03888 | *D. melanogaster* | CycG | OG0003395 | Nurse | Antenna | IIS / TOR | [5] |
| happyhour | DBV15_11653 | *D. melanogaster* | hppy | OG0000352 | Nurse | Antenna | IIS / TOR | [5] |
| insulin receptor isoform (InRL) | DBV15_10224 | *H. saltator* | LOC105183944 | OG0001487 | Nurse | Antenna | IIS | [3] |
| Pi3K21B | DBV15_03081 | *D. melanogaster* | Pi3K21B | OG0002238 | Nurse | Antenna | IIS | [5] |
| Ras oncogene at 64B | DBV15_02802 | *D. melanogaster* | Ras64B | OG0007075 | Nurse | Antenna | IIS / TOR | [5] |
| sarcoplasmic calcium-binding protein 2 | DBV15_05072 | *D. melanogaster* | Scp2 | OG0005773 | Nurse | Antenna | JH: metabolism | [5] |
| shaggy | DBV15_09601 | *D. melanogaster* | sgg | OG0000404 | Nurse | Antenna | IIS / TOR | [3,5] |
| allatostatin A | LOC112454443 | *D. melanogaster* | AstA | OG0007104 | Forager | Brain | JH: regulation of JH biosynthesis | [5] |
| insulin-like growth factor I (IGF1) | LOC112454447 | *H. saltator* | LOC105186969 | OG0004678 | Forager | Brain/Antenna | IIS | [3,4] |
| 5-hydroxytryptamine (serotonin) receptor 2A | DBV15_11483 | *D. melanogaster* | 5-HT2A | OG0000533 | Forager | Antenna | bioagenic amines | [6] |
| octopamine receptor in mushroom bodies | LOC112465659 | *D. melanogaster* | Oamb | OG0001501 | Forager | Antenna | bioagenic amines | [6] |
| tyramine beta hydroxylase | DBV15_00422 | *D. melanogaster* | Tbh | OG0006646 | Forager | Antenna | bioagenic amines | [6] |

| | | | | | | | | |
|---|---|---|---|---|---|---|---|---|
| density regulated protein | DBV15_10283 | *D. melanogaster* | DENR | OG0004331 | Forager | Antenna | IIS / TOR | [5] |
| dreadlocks | DBV15_03530 | *D. melanogaster* | dock | OG0001746 | Forager | Antenna | IIS / TOR | [5] |
| juvenile hormone-inducible protein 26 | DBV15_02220 | *D. melanogaster* | JhI-26 | OG0000832 | Forager | Antenna | JH: regulation of JH biosynthesis | [5] |
| Krueppel homolog 1-like | DBV15_06330 | *H. saltator* | LOC112589664 | OG0008125 | Forager | Antenna | JH: signalling | [3,4] |
| rapamycin-insensitive companion of Tor | DBV15_07973 | *D. melanogaster* | rictor | OG0003860 | Forager | Antenna | IIS / TOR | [5] |
| SAPK-interacting protein 1 | DBV15_07343 | *D. melanogaster* | Sin1 | OG0005622 | Forager | Antenna | IIS / TOR | [5] |
