## Supplemental Table S2 for "Task-specific patterns of odorant receptor expression in worker antennae indicates a sensory filter regulating division of labor in ants"

**Table S2. List of enriched GO biological process terms of the DEG in brain and antenna.**

| Gene Set (DEG) | Tissue | GO ID | Term | Annotated | Significant | Expected | Fisher (adjusted *p*-value) |
|---|---|---|---|---|---|---|---|
| Nurse | Brain | GO:0006412 | translation | 73 | 19 | 1.79 | 0.000 |
| Nurse | Brain | GO:0006879 | cellular iron ion homeostasis | 5 | 2 | 0.12 | 0.006 |
| Nurse | Brain | GO:0009056 | catabolic process | 155 | 6 | 3.79 | 0.024 |
| Nurse | Brain | GO:0030155 | regulation of cell adhesion | 1 | 1 | 0.02 | 0.025 |
| Nurse | Brain | GO:0010040 | response to iron (II) ion | 1 | 1 | 0.02 | 0.025 |
| Nurse | Brain | GO:0030245 | cellulose catabolic process | 1 | 1 | 0.02 | 0.025 |
| Nurse | Brain | GO:0009204 | deoxyribonucleoside triphosphate catabolic process | 1 | 1 | 0.02 | 0.025 |
| Nurse | Brain | GO:0006559 | L-phenylalanine catabolic process | 2 | 1 | 0.05 | 0.048 |
| Nurse | Brain | GO:0006572 | tyrosine catabolic process | 2 | 1 | 0.05 | 0.048 |
| Nurse | Brain | GO:0007160 | cell-matrix adhesion | 2 | 1 | 0.05 | 0.048 |
| Nurse | Brain | GO:0006546 | glycine catabolic process | 2 | 1 | 0.05 | 0.048 |
| Nurse | Brain | GO:0019236 | response to pheromone | 2 | 1 | 0.05 | 0.048 |
| Nurse | Brain | GO:0030334 | regulation of cell migration | 2 | 1 | 0.05 | 0.048 |
| Forager | Brain | GO:0006518 | peptide metabolic process | 89 | 4 | 0.96 | 0.001 |

| | | | | | | | |
|---|---|---|---|---|---|---|---|
| Forager | Brain | GO:0006431 | methionyl-tRNA aminoacylation | 1 | 1 | 0.01 | 0.011 |
| Forager | Brain | GO:0032481 | positive regulation of type I interferon production | 1 | 1 | 0.01 | 0.011 |
| Forager | Brain | GO:0045454 | cell redox homeostasis | 2 | 1 | 0.02 | 0.022 |
| Forager | Brain | GO:0046855 | inositol phosphate dephosphorylation | 4 | 1 | 0.04 | 0.043 |
| Nurse | Antenna | GO:0006412 | translation | 73 | 22 | 8.84 | 0.000 |
| Nurse | Antenna | GO:0006897 | endocytosis | 15 | 6 | 1.82 | 0.006 |
| Nurse | Antenna | GO:0006006 | glucose metabolic process | 10 | 4 | 1.21 | 0.006 |
| Nurse | Antenna | GO:0051726 | regulation of cell cycle | 9 | 3 | 1.09 | 0.014 |
| Nurse | Antenna | GO:0015012 | heparan sulfate proteoglycan biosynthetic process | 3 | 2 | 0.36 | 0.040 |
| Nurse | Antenna | GO:0006886 | intracellular protein transport | 63 | 13 | 7.63 | 0.047 |
| Forager | Antenna | GO:0046854 | phosphatidylinositol phosphorylation | 6 | 4 | 0.57 | 0.001 |
| Forager | Antenna | GO:0046855 | inositol phosphate dephosphorylation | 4 | 3 | 0.38 | 0.003 |
| Forager | Antenna | GO:0005975 | carbohydrate metabolic process | 81 | 17 | 7.74 | 0.004 |
| Forager | Antenna | GO:0045087 | innate immune response | 2 | 2 | 0.19 | 0.009 |
| Forager | Antenna | GO:0009166 | nucleotide catabolic process | 4 | 3 | 0.38 | 0.025 |
| Forager | Antenna | GO:0007229 | integrin-mediated signaling pathway | 9 | 3 | 0.86 | 0.045 |

| | | | | | | | |
|---|---|---|---|---|---|---|---|
| Forager | Antenna | GO:0006166 | purine ribonucleoside salvage | 4 | 2 | 0.38 | 0.047 |
