## Supplemental Table S3 for "Task-specific patterns of odorant receptor expression in worker antennae indicates a sensory filter regulating division of labor in ants"

**Table S3. Behaviors and positions annotations during nest scans.** Nursing was defined as the number of observations a worker antennating, grooming, feeding or carrying a brood item, and its position inside the nest (on the brood or near to the brood pile). Foraging was defined as the number of observations an individual was found outside the nest, collecting food or water.

| **Position** | |
|---|---|
| Inside the nest | On the brood pile |
| | Near the brood pile |
| | Surroundings of the nest |
| | Near the entrance |
| Outside the nest | Chamber 1 |
| | Chamber 2 |
| | Chamber 3 |
| **Behavior** | |
| Brood care | Antennation, grooming, feeding or carrying the brood |
| Nestmate care | Antennation, grooming, feeding or carrying a nestmate |
| Forager | Collecting food/drinking water outside the nest |
| Others | Resting |
| | Walking |
| | Being groomed or fed |
| | Grooming itself |
